# Neurophysiology of mismatch negativity generation: a biophysical modeling study

**DOI:** 10.1101/2025.11.08.687373

**Authors:** Carolina Fernandez Pujol, Joshua Bruce, Ryan V. Thorpe, Stephanie R. Jones, Andrew R. Dykstra

**Author notes:** For correspondence: Carolina Fernandez Pujol. Email addresses (CFP), (JB), (RVT), (SRJ), (ARD).

## Abstract

The mismatch negativity, or MMN, is a ubiquitous evoked brain response elicited by any discriminable change of an otherwise regular stimulus sequence. Despite its potential clinical relevance - the MMN is known to be affected by brain state, lesions, and neurologic/psychiatric disorders - and a growing body of animal work, the underlying neurophysiology of the MMN is not well understood. This hinders translation of circuit-level animal findings and mitigates the utility of the MMN as a neurologic/psychiatric biomarker. Here, we used biophysical modeling to examine the neurophysiological basis of the MMN as measured in a canonical auditory oddball paradigm with frequency deviants (i.e., tones whose frequency was shifted slightly with respect to standard tones). The response to standards was successfully modeled by a typical feedforward-followed-by-feedback input sequence. The response to deviants required additional, prolonged input to supragranular layers, consistent with input from the non-lemniscal thalamus. This additional input resulted in downward-going pyramidal-neuron currents in both layer 2/3 (via indirect somatic inhibition) *and*, critically, layer 5 (via direct apical excitation), which together generated the MMN. The results suggest that current circuit-level models of MMN generation derived from animal models are incomplete, and that further work is required to characterize the underlying neurophysiology of the MMN.

**SIGNIFICANCE STATEMENT:** The mismatch negativity, or MMN, is a change-related brain response and one of the most widely studied brain responses in neuroscience. However, despite its prevalence and potential utility as a biomarker for neurological and psychiatric disorders, its underlying neurophysiology is not well understood. We combined MEG with biophysical modeling to better understand the cells, circuits, and cortical laminae that contribute to deviance processing and the MMN in humans.

## INTRODUCTION

The mismatch negativity, or MMN, is a change-related component of the auditory evoked response (Näätänen et al., 1978) - with counterparts in vision (Pazo-Alvarez et al., 2003; Kimura et al., 2011), touch (Kekoni et al., 1997; Shinozaki et al., 1998), and olfaction (Krauel et al., 1999; Sabri et al., 2005) - and a ubiquitous brain response in neuroscience. Elicited in response to any discriminable violation of an otherwise regular stimulus sequence, the MMN in oddball paradigms (Squires et al., 1975) appears as a difference wave between repeated (i.e., standard) and rare (i.e., deviant) tones (Fig 1).

**Figure 1.**
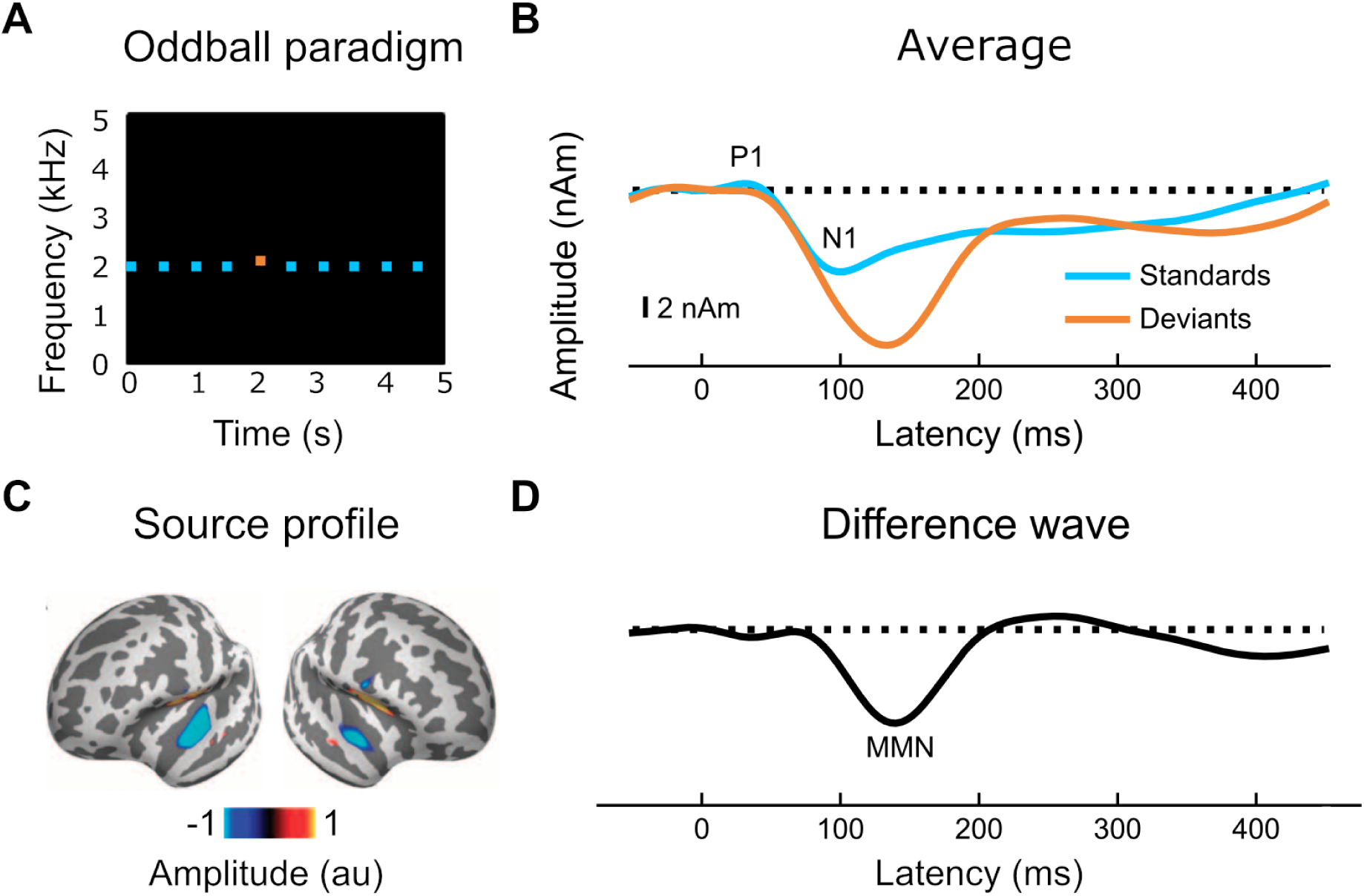
Stimulation paradigm and evoked responses. (A) Sample trial consisting of a sequence of standard tones (blue squares) and a single deviant tone (the oddball) of higher frequency in position five (orange square). (B) Average of standards and deviants across hemispheres. (C) Source profiles between 100 and 150 ms for deviants minus standards. (D). Difference wave between the recorded response to the standard and the recorded response to deviant tones. Adapted from Dykstra and Gutschalk, 2015. Negative is down.

Functionally, the MMN is thought to reflect automatic change detection, yielding insight into how the brain processes mismatches between expected and actual sensory input (Garrido et al., 2009). Due to this interpretation, and perhaps also the ease with which it is measured, many studies have investigated the utility of the MMN as a biomarker in neurological and psychiatric disorders. However, though MMN generation does not require task engagement with deviant stimuli, it may require conscious perception of the contextual regularity (Sussman et al., 2014; Dykstra and Gutschalk, 2015), consistent with the fact that both prefrontal lesions and attention-related cognitive impairments reduce or abolish the MMN (Alho et al., 1994; Näätänen et al., 2011, 2014).

Anatomically, the MMN arises from auditory cortex (AC) on the anterolateral superior temporal plan (Näätänen and Alho, 1995; Dykstra and Gutschalk, 2015), with a possible later, secondary source in inferior frontal cortex (Näätänen and Alho, 1995). However, despite increasing work in experimental animals and pharmacological studies in humans, the neurophysiology underlying the MMN is not well understood. MMN generation has been linked to both *N*-methyl-D-aspartate (NMDA) receptor function and somatostatin (SST) driven disinhibition of pyramidal neurons (PNs). NMDA receptor antagonists (ketamine, e.g.) attenuate the MMN (Javitt et al., 1996; Javitt, 2000; Heekeren et al., 2008; Näätänen et al., 2011; Rosburg and Kreitschmann-Andermahr, 2016) while NMDA potentiators increase it (Lavoie et al., 2008). Reduced SST activity decreases MMN-like responses (Hashimoto et al., 2008; Hamm and Yuste, 2016), while enhanced SST activity disinhibits PNs (Ma et al., 2010) via inhibiting parvalbumin (PV) interneurons (Cottam et al., 2013).

Taken together, these findings have inspired several models of MMN generation, which differ regarding which neurons, circuits, and layers are most involved (Fishman and Steinschneider, 2012; Lakatos et al., 2020; Ross and Hamm, 2020; Varela et al., 2024). While most models include populations of layer 2/3 (L2/3) PNs (and associated SST and PV interneurons) that are differentially and selectively tuned to standard and deviant stimuli (Ross and Hamm, 2020), the extent to which deeper layers (L5/L6), downstream effects, and different thalamic subdivisions are involved is less understood. For example, macaque recordings show that while the lemniscal pathway does not show deviance encoding (Lakatos et al., 2020) (including ventral medial geniculate body, or MGBv, and its primary target, L4 of primary auditory cortex), the non-lemniscal pathway does (including the dorsal and medial subdivisions of the MGB and L1 of cortex, which, along with L5, receives projections from MGBd/m).

There are also discrepancies in recording sites, types, and latencies at which MMN-related responses are found. While the auditory MMN in humans arises from *non-primary* auditory cortex at ∼150 ms, most animal recordings are from *primary* sensory areas and show much earlier deviance responses. Even *within* animal studies, exactly what is measured varies between laminar probes, L2/3 recordings, or two-photon microscopy. To address these discrepancies and bridge between MMN work in humans and animal models, we used biophysical modeling to model how the auditory MMN might arise from specific neurons, circuits, and layers. We took into account data and predictions from both rodent and primate data to examine hypotheses of MMN generation. Modeled peaks include the auditory P1-N1 complex (seen in response to both standards and deviants) and the MMN, which includes a larger and more prolonged N1 for deviants.

## METHODS

### Data

All data used for this modeling study came from prior work that examined MEG responses to target-tone streams embedded in random, multitone maskers (Dykstra and Gutschalk, 2015). The responses we modeled here came from the control blocks which consisted of a standard oddball paradigm. Details of the experimental design, acoustic stimulation, task, and data analysis can be found there. We provide brief overviews below.

### Auditory stimulation

The paradigm used to acquire the MMN signal was a standard oddball paradigm consisting of a stream of nine repeating pure tones (the standards) and a single deviant - occurring at one of positions 3-8 in the 9-tone sequence - which differed only in frequency from the standard tones by 1/12 of an octave in either direction in a completely counterbalanced manner. Each sequence was 5 seconds long, each tone was 100 ms long, and the stimulus-onset asynchrony between successive tones was 500 ms). All tones were presented at comfortable listening levels well above sensory threshold. The participants (N=20) were asked to passively listen to the stream of repeating tones. All responses to the standards and deviants were combined to obtain average evoked related responses for standards and deviants per hemisphere. For the purposes of this modeling study, the responses were also averaged across hemispheres.

### MEG acquisition and analysis

Details about MEG acquisition, preprocessing, and analysis can be found in earlier work (Dykstra and Gutschalk, 2015; Fernandez Pujol et al., 2023). Briefly, the MEG responses were acquired using a Neuromag 122 system with orthogonal pairs of planar gradiometers at each of 61 locations around the head. MEG was acquired at a sampling rate of 500 Hz with a 160-Hz online lowpass filter. The data were then bandpass-filtered (0.5–15 Hz) offline, and a PCA-based artifact-rejection algorithm was used to remove eye blinks and saccades. The data was then epoched from -50-450 ms around the onset of the target tones and binned into one of two categories: standard target tones and deviant tones.

HNN requires M/EEG responses to be source-localized and the orientation of the dipole current in relationship to the cortical surface to be estimated. The data used for the current study were source-localized using a combination of anatomically constrained, noise-normalized minimum norm estimates (Dale and Sereno, 1993; Hämäläinen and Ilmoniemi, 1994) also known as dynamic statistical parametric mapping (Dale et al., 2000) and sparse inverse estimates based on an L1/L2 mixed norms (Gramfort et al., 2012). Individualized structural MRI scans, including T1-weighted magnetization-prepared rapid gradient echo (MPRAGE) and multiecho fast low-angle shot (FLASH) sequences, were acquired for each subject using a 3-T Siemens TIM Trio scanner. Individual source spaces and boundary element models were created, and the source estimates were obtained for each subject’s left and right AC. This was all carried out using MNE-Python and Freesurfer (Fischl, 2012). Alignment of the MRI and MEG coordinate frames was done in MNE-Python (Besl and McKay, 1992). After source localization in individual subjects, the grand average of both hemispheres was calculated and modeled for each response (standard vs. deviant tones. These source-localized signals (units of ampere-meters, Am) correspond to the net longitudinal current flow within postsynaptic PNs—also known as primary currents—and can be thought of as primary current dipoles (with a defined strength and direction). HNN works by simulating the primary currents given a set of inputs and corresponding parameters and outputs the primary current dipoles which can then be directly compared to the empirical, primary current dipoles.

### Model

The Human Neocortical Neurosolver (HNN) (Neymotin et al., 2020) is designed to model source-localized activity from a single cortical column to develop and test hypotheses about the circuitry giving rise to event-related responses of interest and has been previously applied to auditory-evoked responses associated with both conscious perception of tones in noise (Fernandez Pujol et al., 2023) and hemispheric differences / contralateral dominance (Kohl et al., 2021).

HNN provides a canonical neocortical column consisting of L2/3 and L5 (the layers with neurons whose dendrites contribute most to activity recorded with MEG outside of the head) (Fig. 2) (Neymotin et al., 2020; Jas et al., 2023). Each layer contains both PNs (modeled as multiple compartment neurons with one oblique, two basal, and one apical dendrite) and inhibitory basket cells (modeled as single-compartment neurons). The basket cells represent fast-spiking GABAergic interneurons. The somata of the L2/3 neurons and the somata of the L5 neurons reside in L2/3 and L5 respectively, with the apical dendrites of the PNs spanning from their respective layers to the cortical surface. L4 (i.e. the granular layer) is not modeled and presumed only to relay information to the supra- and infra-granular layers (at least insofar as generation of M/EEG signals is concerned). Each layer consists of a ten-by-ten grid of PNs with basket cells added in a 3:1 PN-to-basket-cell ratio for a total of 270 cells (full model not shown). A net current dipole signal is simulated by calculating the intracellular current flow within the PN dendrites multiplied by the length of the dendrites, and summing across all PNs in both L2/3 and L5. The result is a signal in nano-ampere-meter (nAm) units and allows for a one-to-one comparison between the simulated and sourced localized M/EEG data in the same units. Although this number of neurons in the default HNN model is fixed, a scaling factor is applied to the modeled data to account for the overall amplitude of the M/EEG response to be modeled and provides an estimate of the number of neurons contributing to the recorded data. Here, the scaling factor is 450.

**Figure 2.**
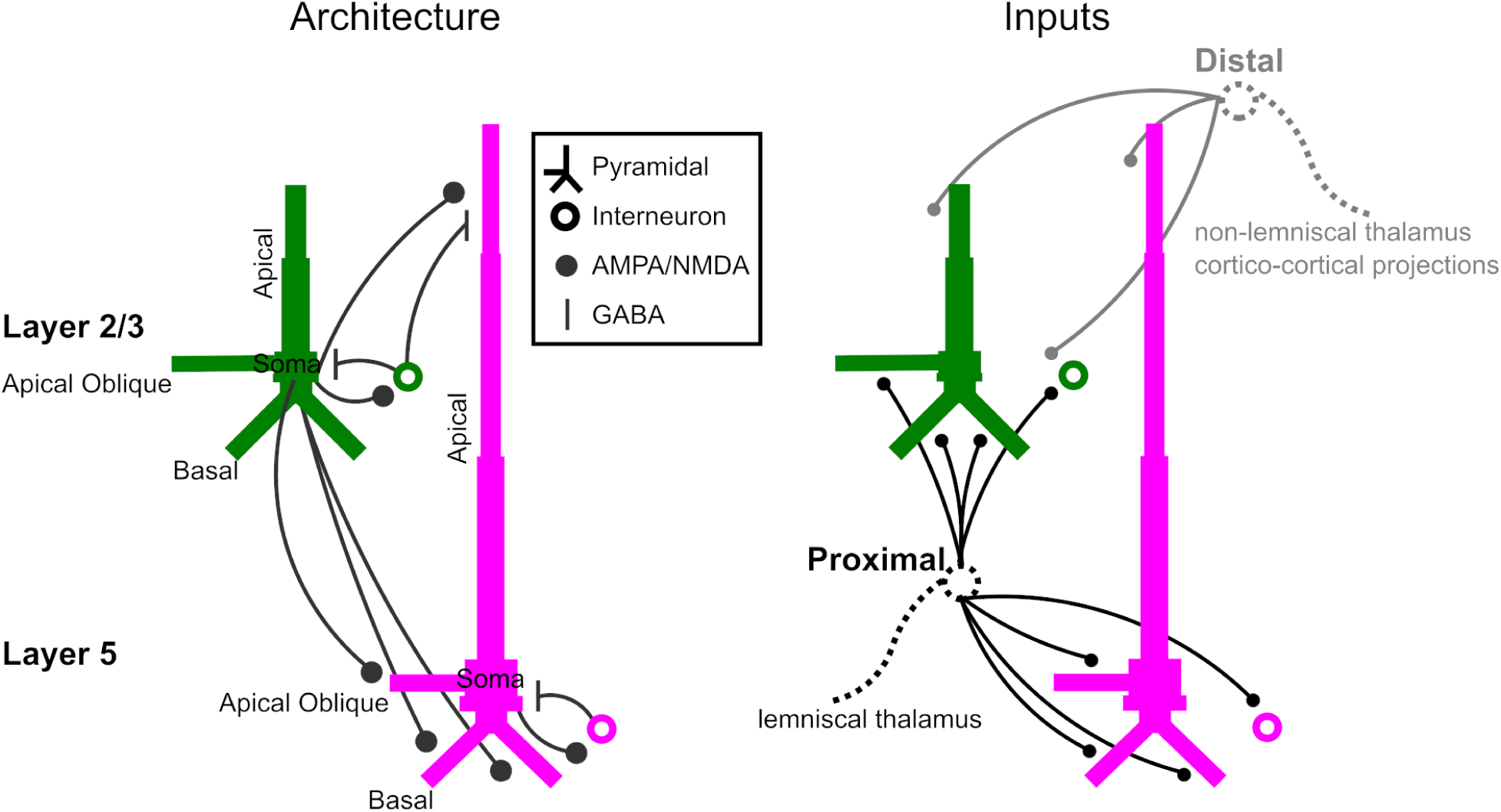
The Human Neocortical Neurosolver. Architecture / underlying connectivity of HNN model cortical column and inputs (proximal input in black and distal input in gray).

The PNs are connected with each other and also connected to the basket cells via excitatory (AMPA/NMDA) synapses (Fig. 2, black circles). The basket cells are connected to the PNs via fast and slow inhibitory (GABAA/GABAB) synapses (Fig. 2, black lines). In the default HNN model used here, the strengths/weights of these local connections (in μS) were fixed to reflect the dynamics of a canonical neocortical column, but can be modified with sufficient physiological justification (for example, in non-canonical contexts such as disease models). HNN uses standard Hodgkin-Huxley conductance equations to describe the biophysical characteristics of neuronal cell membranes, and computes current flow between neuronal compartments using cable theory.

At rest, the network does not generate any activity and instead must be driven with exogenous inputs to model the experimental conditions of interest. Here we consider two kinds. Proximal inputs (proximal to the perisomatic compartment) arrive from the lemniscal thalamus and synapse onto the basal and oblique dendrites of the L2/3 and L5 PNs and onto the basket cells. They reflect the main pathway through which sensory information reaches the primary sensory cortices. Because they are excitatory, proximal inputs cause glutamate to bind to AMPA/NMDA receptors on the postsynaptic cell, which in turn causes Na^+^ ions to enter (and few K^+^ ions to leave) the cell. The result is net positive influx of charge at the perisomatic compartment of the postsynaptic neuron. This current brings about a local, extracellular negativity at the perisomatic compartment, rendering the extracellular space near the apical compartment more positive. Due to the potential difference between the apical and perisomatic compartments, intracellular current flows longitudinally and upwards. The result is a current dipole with a positive peak. Additionally, if the excitatory proximal input elicits a spike in the PN, this induces a large back-propagation of current up the PN dendrites. Distal (to the perisomatic compartment) inputs arrive from the non-lemniscal thalamus or higher-order cortical areas and synapse onto the apical dendrites of the L2/3 and L5 PNs and onto the L2/3 basket cells. They reflect input from either the non-lemniscal thalamus (which is often modulatory and thus may be involved in processes that require attention (Harris and Mrsic-Flogel, 2013; Harris and Shepherd, 2015)) or from other cortical areas. The result of driving the model column with a distal input is net positive influx of charge at the apical compartment of the postsynaptic neuron which brings about a local, extracellular negativity at the apical compartment, creating longitudinal intracellular currents within the PN dendrites that flows downward. The result is a current dipole with a negative peak. Additionally, if the excitatory distal input generates a dendritic calcium spike, this also induces a downward current. The proximal and distal inputs can also elicit action potentials in the basket cells, which provide inhibition to either the somal or distal dendrites of the PNs (Fig. 2). Inhibitory currents onto the soma will pull current flow down the dendrites, and on the distal dendrites it will pull current flow up. In total, the observed current dipole signal comes from a combination of the direct effect of the exogenous input onto the PN dendrites and the resulting induced local network dynamics. By examining the current dipole simultaneously with spiking activity in the various cell types in the model, one develops a complete picture of how the underlying multi-scale network activity produces the observed macroscale current dipole signal.

The networks from which proximal and distal inputs arrive (lemniscal thalamus, non-lemniscal thalamus, and higher-level cortical areas) are not explicitly modeled but their outputs are simulated by input spikes (action potentials) that drive the postsynaptic neurons. Each input comes with a set of parameters that can be adjusted accordingly and include: the average start time for the spike (units of ms), the standard deviation of the average start time (units of ms), the strength of the input on the postsynaptic cell (units of μS), the time before the postsynaptic cell receives the spike (ms), and the number of spikes. Stochasticity in the response across trials comes from jitter in the standard deviation of the input time on any given trial. Here we simulated N=10 trials (thin gray curves) and averaged across trials (thick colored curves).

Here, we used HNN to model the MMN, namely the larger negative response between 100-200ms (Fig. 2). The MMN, which arises from AC, is thought to represent an automatic/involuntary attentional switch to a change in the acoustical environment; a biologically vital warning response to change.

### Hypothesis testing with the HNN model

The process of using HNN to make detailed predictions on the mechanisms of generation of recorded data begins by starting with the distributed pre-tuned HNN model and adjusting parameters of the exogenous input (and/or local network features; not examined here) to test hypotheses on the origin of the data at hand. Parameter adjustments are performed by first hand-tuning, and then applying automated parameter estimation methods in HNN (Bergstra et al., 2011; Jas et al., 2023). To model the response to the standard tones, we began with a proximal-distal input sequence used in our previously published model of a typical auditory evoked response (Kohl et al., 2021; Fernandez Pujol et al., 2023) and re-tuned the model to fit the standard responses in the data studied here.

To model the response to the deviant tones, we kept the first two inputs fixed (as in the response to the standard tones), and introduced a second distal input.

It is important to note here that in the frequency oddball paradigm, due to tonotopic organization of the auditory cortex, the response to deviants likely arises primarily from a cortical column that is distinct from that in which the response to standards arise. This distinction was accounted for, however, as the trials were counterbalanced such that the number of deviants (and standards) occurring at either the lower or higher frequencies were matched. Furthermore, the MMN observed in frequency oddball paradigms is known to arise from areas of auditory association cortex that are anterolateral to A1, as was also found in our data (cf. Fig. 1C) (note, also, that we did not find evidence of a frontal MMN sub-component, and therefore focused exclusively on the component arising from AC). As such, the net current dipole estimate in our data represents the forward solution from this area which, contrary to primary auditory cortex, is less likely to be tonotopically organized. In HNN, we are simulating activity in a single canonical cortical model and using it to represent two experimental conditions with responses that are possibly spatially distinct (i.e., standard- and deviant-tone responses). We use the model to predict the response to standard and deviant stimulus separately, assuming that only one cortical area (i.e. the one primarily responding to either the standard or the deviant) is active at given time. Both responses were captured within the region of interest defined during source estimation. However, given the nature of the source estimation approach, it is important to note that the localized activity may include crosstalk from other brain regions, that is, signals originating elsewhere may partially contaminate the estimated activity of the target region. For our purposes, this implies that a single model can effectively represent the circuit dynamics underlying both the standard and deviant conditions.

## RESULTS

### Overall Biophysical Model Configuration

Details of the biophysical model column implemented by HNN can be found in the Methods section and have also been described elsewhere (Jones et al., 2007; Neymotin et al., 2020; Fernandez Pujol et al., 2023). Briefly, HNN models the primary currents contributing to MEG signals with a reduced cortical column circuit that includes two layers (L2/3 and L5) and cell types (PNs & inhibitory interneurons in the form of basket cells). The model is activated (i.e. driven) via two types of synaptic input: proximal or distal. Proximal inputs (Fig. 2, black) target the basal and oblique dendrites of PNs and the soma of basket cells and are thought to reflect feedforward information from either the lemniscal thalamus or an earlier cortical area. This influx of ions generates a dipole that drives current up within the PN apical dendrites, and can generate somatic action potentials that backpropagate up dendrites, towards the cortical surface. Evoked inhibition onto the distal dendrites will also pull current flow up. Distal inputs (Fig. 2, gray) target the apical dendrites of PNs and the soma of L2/3 basket cells and reflect projections from either the non-lemniscal thalamus or higher-order cortical areas. This influx of ions together with potential dendritic calcium spikes generates a dipole that drives current down the PN dendrites, towards the white matter. Evoked somatic inhibition will also pull current flow down.

The primary model output from an HNN that is compared to data is the net dipole moment from the cortical column, and is calculated from the net intracellular currents in the PN dendrites, in the same units (nAm) as the source-localized M/EEG signal that is being modeled. HNN’s outputs also include individual layer activity (laminar profiles), spiking activity of individual cells in the network, and local field potentials (LFPs). The laminar profiles estimate the individual contributions of L2/3 and L5 PN dipoles to the net current dipoles (sum of activity in L2/3 and L5). The individual cell spiking activity plots describe how the microcircuit activity in the column might give rise to the aggregate macroscale current dipole signal. The LFPs estimate the extracellular potentials as measured across the depth of the model cortical column via a simulated linear multielectrode array (albeit without a granular layer in the model). Current source density, or CSD, can also be estimated as the second spatial derivative of the LFP, and is an estimate of the current sources that generate the simulated extracellular potentials. Collectively, these allow us to study the neurophysiology of the MMN at the level of single cells and circuits.

Simulating Standards and Deviants in the Primary Current Dipole

We first modeled the response to standard tones, which consists of a positivity at around 50 ms and a negativity at around 100 ms (the auditory P1-N1 complex) (Fig. 4A). To reproduce this morphology, we began with the same first two inputs used to model the auditory P1-N1 complex in the context of a multitone masking paradigm for a previous study (Fernandez Pujol et al., 2023). Because the morphology and amplitude of the responses differ from that study, we first hand-tuned the parameters until a good fit was achieved. We then set the excitatory proximal input onto the L2/3 basket cells to a weight of 0 μS based on the fact that in the adapted state circuit model proposed previously (Ross and Hamm, 2020), the basket cells (i.e. PV interneurons) are disinhibited. The parameters in the model were then optimized to fit the standard response using HNN’s optimization tool. For the distal input, the NMDA and AMPA conductances onto L2/3 interneurons were fixed at 0 μS, and the maximum possible value for the NMDA conductances onto the L2/3 PNs was set to not exceed 1 μS. This low NMDA conductance value onto L2/3 PNs is representative of synaptic depression that arises when the PN population selective for the standard-frequency are repetitively stimulated (i.e., adaptation). This resulted in an R^2^ value of 0.59 and a root mean-squared error (RMSE) value of 3.43 nAm (see Fig. 3 for all parameter values). The final, optimized model contains two driving inputs - a proximal (i.e., feedforward) input at 49.8 ms and a distal (i.e., feedback) input at 91.6 ms - and produced a good fit to the data up through the N1 response [R^2^ = 0.59, root mean-squared error (RMSE) = 3.43] (Fig. 4A). We did not attempt to model the slow return to baseline for either standards or deviants, as such the model and data diverge at the later time points.

**Figure 3.**
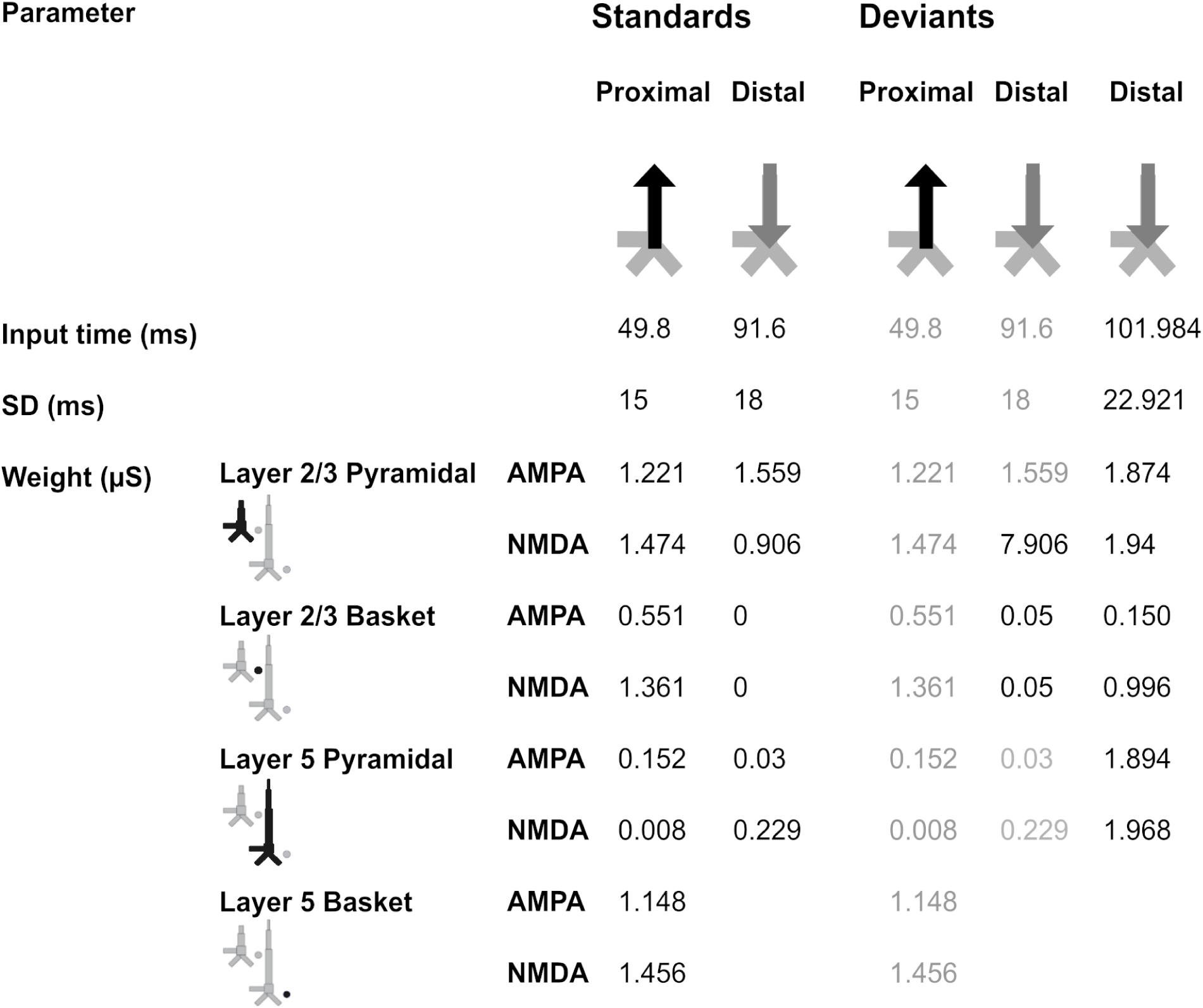
Driving inputs (and parameters) used to model the responses to the standard and deviant tones. The model column used was adapted from HNN’s original model (Jones et al., 2009), to include a more biologically accurate distribution of Ca^2+^ channels on L5 PNs, as in (Kohl et al., 2021; Fernandez Pujol et al., 2023).

**Figure 4.**
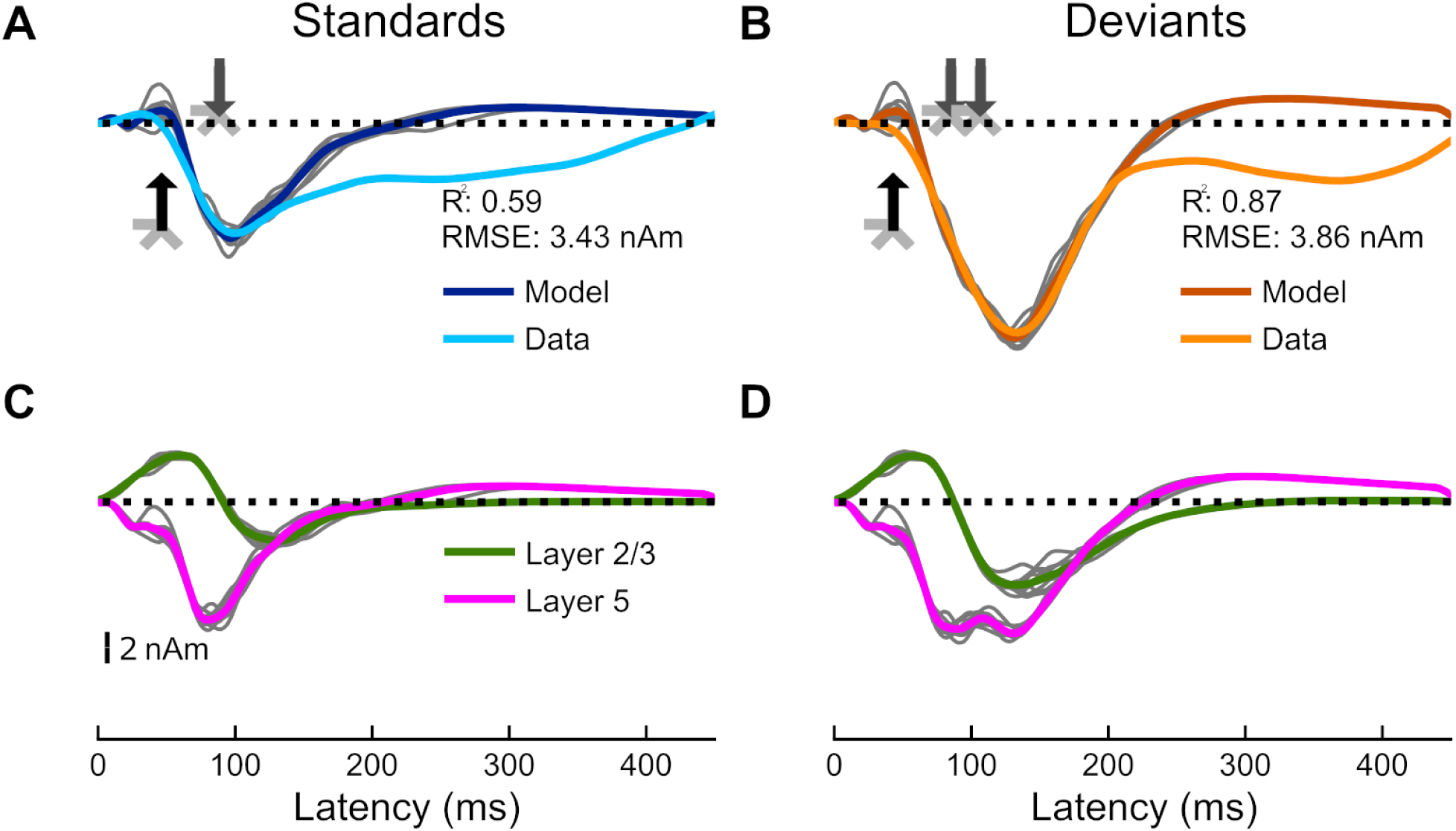
Current dipoles and laminar profiles. (A) HNN simulation for the average response to the standard tones (dark blue trace). Input spikes are sampled from a Gaussian distribution on each trial for a total of 10 trials (gray traces). A proximal input (49.8 ms) followed by a distal input (91.6 ms) drive the network. R^2^ between empirical (light blue trace) and simulated data (dark blue) is 0.59 and RMSE is 3.43 nAm. (B) HNN simulation for the average response to deviant tones (dark orange traces). The proximal and distal inputs used to model the response to standard tones (with a few changes), and an additional distal input (101.98 ms), drive the network. R^2^ between empirical (light orange trace) and simulated data (dark orange) is 0.87 and RMSE is 3.86 nAm. (C-D) Laminar profiles for the responses to the standard and deviant tones. The contribution of L2/3 (which reflects the longitudinal currents in layer-2/3 (L2/3) PNs only) is plotted in green and that of layer 5 (which reflects the longitudinal currents in L5 PNs only) in magenta. Note that the model traces in panels A and B reflect the aggregate of the green and magenta traces in panels C and D. The corresponding input parameter values are displayed in Fig. 3. The network from which the simulated dipole activity arises consists of 90,000 cells.

To simulate the response to the deviant tones (Fig. 4B), which contains an additional negativity between 100 and 180 ms (i.e., the MMN), we initially explored modifications to the input parameters within the standard model. Starting with the model tuned to a standard response, we increased the NMDA conductances of the distal input onto the L2/3 PNs to a value of 7.906 μS under the assumption that the PN population selective for the frequency of the deviants does not experience synaptic depression like standards do. We then slightly increased the excitatory distal input onto the L2/3 basket cells from 0 μS to 0.05 μS assuming the PV interneurons do not experience synaptic depression when the PN population selective for the deviants is stimulated. However, solely increasing the NMDA conductances of the L2/3 PNs did not sufficiently increase the size of the additional negativity present for deviant tones and was thus unable to account for the MMN (Fig. S1B).

Given the observed CSD activity in the supragranular layers of macaque AC, indicative of input from either the non-lemniscal thalamus or higher-order cortical regions (Javitt et al., 1996; Fishman and Steinschneider, 2012; Lakatos et al., 2020; Ross and Hamm, 2020), we tested the hypothesis that an additional distal input (101.98 ms) could account for the MMN. A second distal input was simulated and the parameter values were optimized to fit the deviance response (while keeping the parameters of the first two inputs fixed). This final model for the deviant tones better reproduced the amplitude and morphology of the response to deviant tones (and thus the MMN) (R^2^ = 0.87, RMSE = 3.86 nAm) (Fig. 4B).

The laminar-specific contributions to the simulated waveforms (Fig. 4C and D) show that the difference between responses to deviant and standard tones is due to additional negative-going current activity in both L2/3 (magenta traces) and 5 (green traces). All inputs to the model and their corresponding parameters can be found in Fig. 3.

### Corresponding Network Spiking Activity

The network spiking activity reflects the dynamics of the neocortical circuit at the level of individual cells and contributes in often non-intuitive ways to the generation of the net primary current dipole signal. In response to the standard tones (Fig. 5A), the model shows (i) spiking of both the L2/3 and L5 PNs (solid green and magenta triangles), and (ii) sustained basket-cell firing in both L2/3 and L5 (open green and magenta circles). In response to the deviant tones (Fig. 5B), the model shows additional spiking of all cell classes except for L2/3 PNs (Fig. 6). Notably, the largest difference in spiking between standards and deviants occurs in the L2/3 basket cells, with significantly more firing on deviant trials induced from the additional first and second distal input onto these cells (see Fig. 3). This increased firing creates more somatic inhibition on the L2/3 PN pulling current flow down to PN dendrites. Additionally, the second distal input (and significantly stronger first distal input, see Fig. 3) directly onto the L2/3 PNs helps push current flow down the dendrites via synaptic and dendritic calcium dynamics. Together, these downward currents create a larger and longer duration negative primary current response in L2/3 on deviant trials (compare green curves in Fig. 2 and 3). The primary current response in L5 is of longer duration on deviant trials with a second negative peak appearing just after the second distal input, however the magnitude remains fairly consistent (compare magenta curves in Fig. 4 C and D). This second peak emerges from the second distal input pushing additional current flow down the L5 PN dendrites. This input also produces an initial slight increase in firing in the L5 PNs, which in turn drives increased firing in the L2/3 inhibitory neurons that quickly stops the increased PN firing and continues to pull the current downward. The resulting net primary current dipole is the sum of the L2/3 and L5 dipole responses. Notably, our model deviates from the data after the N1 peak, and this was not a target of our investigation (cf. Discussion).

**Figure 5.**
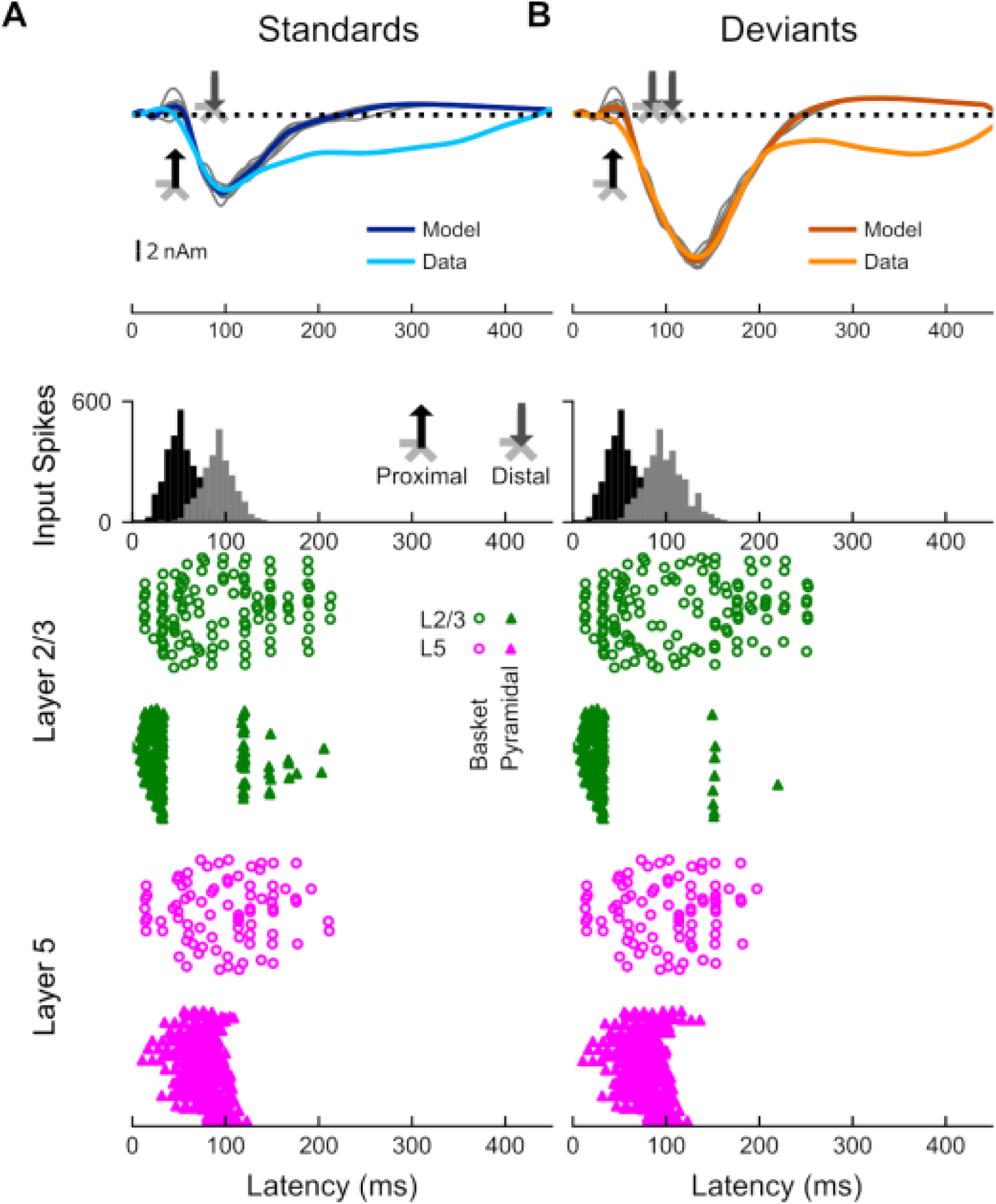
Network spiking activity from a single trial across all neurons of the model. (A) Network spiking activity for the standard tones. (B) Network spiking activity for the deviant tones.

**Figure 6.**
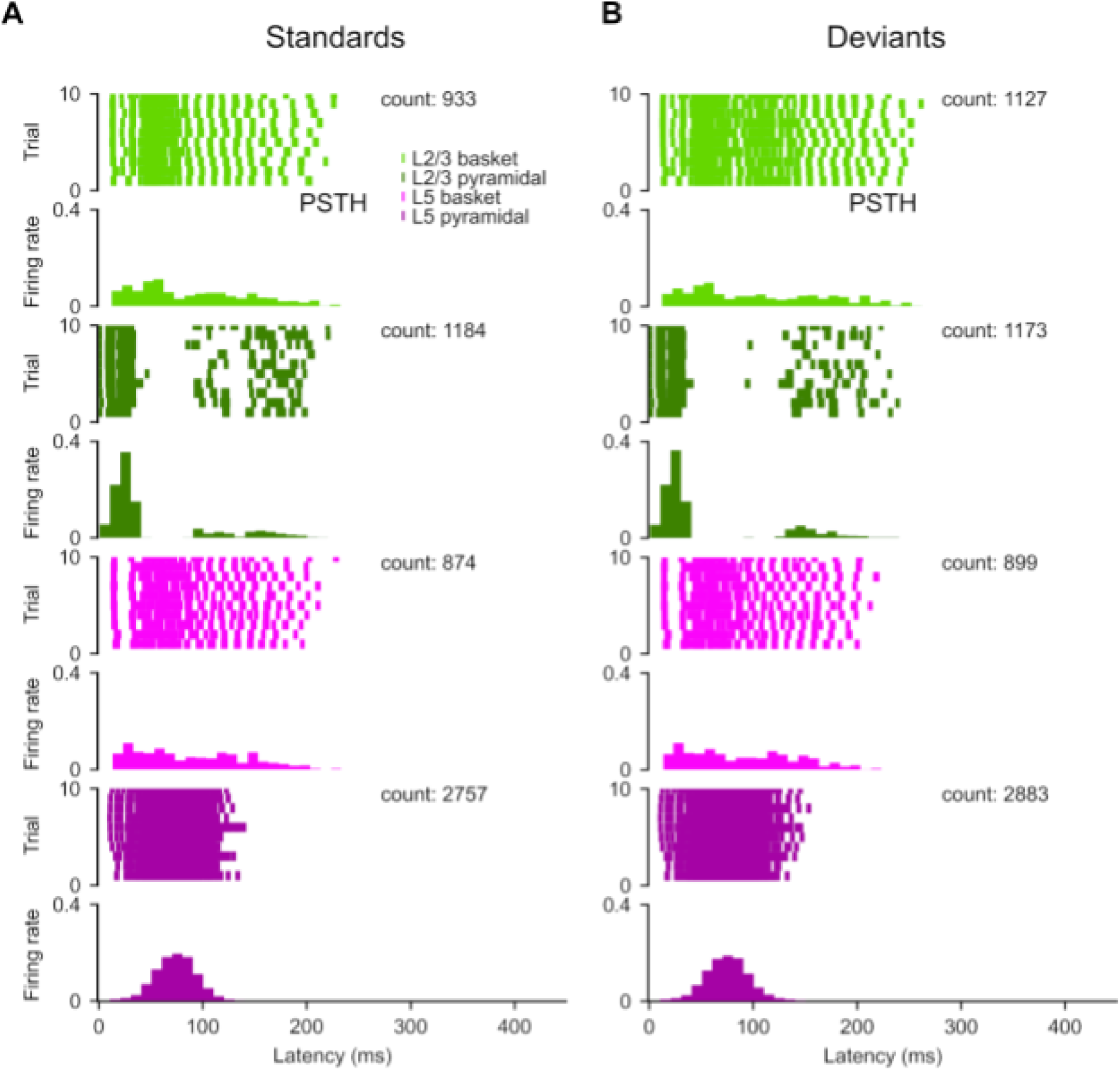
Raster plots and Peri-Stimulus Time Histograms (PSTH) across all neurons and all trials in the network. Each line in a raster plot indicates a spike corresponding to that time point and trial number. The spike times are aggregated into bins, counted, and normalized in the corresponding PSTH. The curves (black) are the kernel density estimate of the distribution.

We also examined LFP/CSD simulations. These revealed deviant-standard differences in CSD sink/source pairs within supragranular and infragranular layers, aligning with findings from animal studies (see Supplementary Materials for description of LFP/CSD simulations). Furthermore, we investigated two alternative models of MMN generation but found them insufficient to accurately capture the observed responses (see Supplementary Materials).

### Simulated Local Field Potentials and Current Source Density

We simulated the layer specific LFP and CSD signals produced by the final standard and deviant simulations using functions distributed with HNN (Neymotin et al., 2020; Jas et al., 2023). These simulations provide further targets for validation of model-based predictions. A simulated linear multi-electrode array (50 electrode contacts) was placed in the center of the model column to measure the extracellular potentials as a function of depth. The length of the hamming window used to smooth the simulated dipole waveform was 30 ms. LFPs reflect integrated transmembrane currents (Buzsáki et al., 2012). Notably, our model does not contain cells in the granular layer (L4), and the LFP/CSD in those layers reflects only the activity emerging from the cells in the other layers; thus predictions from L4 are incomplete. The LFP plots (Fig. 7) show that the response to the standard tones is associated with stronger field activity in the deeper layers (despite the fact that both the proximal and distal input to the L2/3 is stronger (Fig. 3). This aligns with the strong and synchronous somatic spiking that occurs in the L5 PNs during this time window (see Fig. 6). The LFPs for the response to the deviant tones (Fig. 7B) show that the MMN is associated with an increase in LFP in the superficial layers near 150ms, which aligns with more synchronous PN firing in L2/3 in this time window.

**Figure 7.**
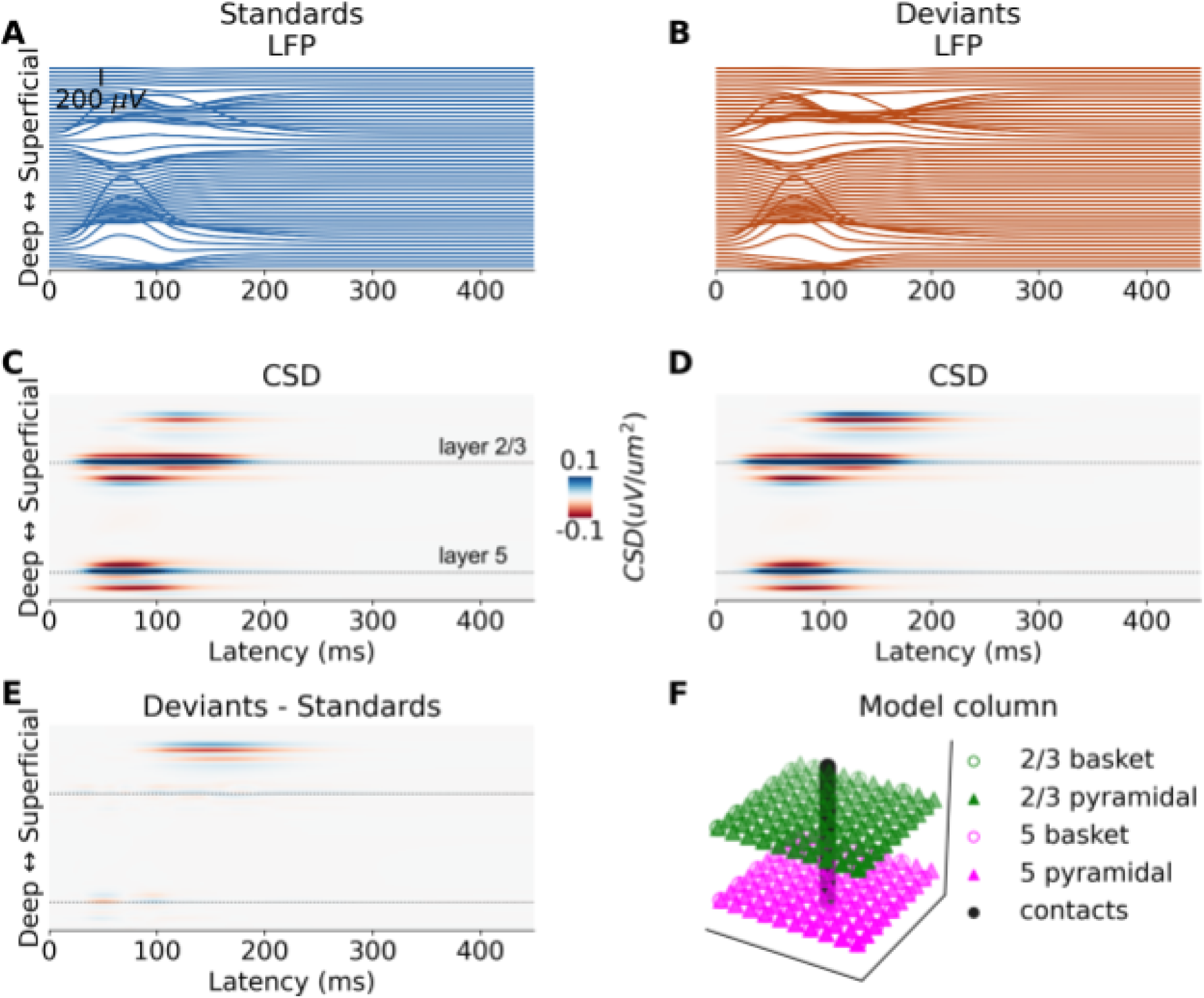
Local Field Potentials and Current Source Density. **(A)** Simulated average LFP for the standard tone responses. The LFP are estimated from a linear multi-electrode array of 50 electrode contacts, **(B)** Simulated average LFP for the deviant tone responses. **(C-D)** Corresponding simulated average CSD, **ļE)** CSD difference between deviants and standards. The model MMN shows a source-sink-source configuration largely in the superficial layers. **(FJ** The model column and simulated 50-contact linear electrode array from which LFPs and CSD were estimated.

The CSD plots are generated by computing the second spatial derivative of the LFPs and provide an estimate of the current sources and sinks that generate the measured LFPs. A negative CSD indicates current flow into the cell (current sink; red) and a positive CSD indicates current flow out of the cell (current source; blue) (Mitzdorf, 1985; Einevoll et al., 2013), and can be interpreted as current flow out and into the extracellular medium, respectively. The CSDs can more precisely localize transmembrane currents than the LFPs since they do not suffer from far-field contamination arising from volume conduction (Kajikawa and Schroeder, 2011; Einevoll et al., 2013; Hindriks et al., 2016). In actual recordings, LFPs receive non-local contributions from up to 1 cm away from their origin (Kajikawa and Schroeder, 2011), which our model does not simulate.

The CSD plots show that the response to the standard tones (Fig. 7C) is mostly associated with a strong source-sink configuration in the infragranular layers. The response to the deviant tones (Fig. 7D) show increased sink/source activity in both the supragranular and infragranular layers. We computed the difference between the CSD of the deviants and the CSD of the standards (Fig. 7). At the latency of the MMN (∼140 ms), the difference of deviant-related synaptic activity is not biased towards either layer and occurred nearly equally in both supra- and infragranular layers; this is distinct from some laminar recordings of the MMN in primary auditory cortex of non-human primates, which show stronger superficial sink/source pairs (Javitt et al., 1996; Fishman and Steinschneider, 2012; Lakatos et al., 2020). This may relate to differences between our reduced model and in-vivo recordings, a point that we return to in the Discussion.

## DISCUSSION

We used biophysical modeling to investigate the neurophysiological mechanisms underlying deviance detection and the MMN in human AC. While both standard and deviant tones elicited strong N1 responses, the P1 was weaker for deviants likely due to temporal overlap with a large negative-going component. Both responses were modeled by a proximal-distal input sequence, which accounted for the P1-N1 complex (Figs 3 and 4) (Kohl et al., 2021; Fernandez Pujol et al., 2023). The MMN was best captured by a combination of increased L2/3 NMDA conductances during the first distal input and a second, delayed distal input to L2/3 and L5 PNs and L2/3 inhibitory neurons. MMN generation involved (i) increased and prolonged spiking in L2/3 inhibitory neurons, producing downward current in L2/3 PN dendrites, and (ii) a strong, sustained current down L5 PN dendrites induced by the second distal input. Importantly, increasing L2/3 NMDA conductances alone was insufficient to reproduce the MMN, highlighting the necessity of a second distal input.

### Model Interpretation

Our results, together with prior macaque data (Fishman and Steinschneider, 2012; Lakatos et al., 2020) and previous predictions (Ross and Hamm, 2020), suggest the following circuit interpretation.

For standards, lemniscal thalamic input arrives to L4 neurons tuned to the standard frequency, where it is relayed to both PV interneurons and the proximal dendrites of PNs in L2/3 and L5. This proximal input (∼50 ms) drives current up the PN dendrites (towards the cortical surface), producing the P1. The first distal input (∼90 ms), targeting apical dendrites of L2/3 and L5 PNs, represents local cortical feedback exhibiting synaptic depression due to repeated stimulation (see Limitations for discussion of stimulus-specific adaptation, or SSA). This was modeled as reduced NMDA conductance in L2/3 PNs and no NMDA/AMPA conductance in L2/3 interneurons. The resulting downward current generated the N1. For deviants, the A1 population tuned to the deviant frequency is unadapted, producing stronger NMDA-mediated distal input to L2/3 neurons, consistent with rodent data (Ross and Hamm, 2020).

Some models propose that deviant-related information is fed back to the lemniscal thalamus (Varela et al., 2024) via L6A corticothalamic (CT) neurons (not explicitly modeled here). When relayed to “burst-ready” thalamocortical neurons (TC), this could generate excitatory feedback to L4, producing a positive-going deflection. While such mechanisms might underlie early deviance-related positivity (Grimm and Escera, 2012), they are unlikely to explain the MMN’s characteristic negativity polarity.

Macaque data implicate the non-lemniscal thalamus and its cortical targets in superficial layers in deviance detection (Javitt et al., 1996; Fishman and Steinschneider, 2012; Lakatos et al., 2020). Laminar recordings show the largest deviant-standard differences in supragranular layers (L1-L2/3) and, to a lesser extent, L5 of A1. Deviant-related current sinks are concentrated in L1, a major target of cortico-cortical and non-lemniscal thalamic projections. Although the precise circuitry is unresolved, L6B neurons, which receive strong excitatory input from L2/3 (Zolnik et al., 2020; Feldmeyer, 2023), provide a potential pathway from cortex to non-lemniscal thalamus (Carbajal and Malmierca, 2018; Zolnik et al., 2020, 2024; Shepherd and Yamawaki, 2021; Feldmeyer, 2023; Whyte et al., 2024). We therefore propose that cortical prediction (memory trace) is compared to sensory input in a subcortical circuit, with the outcome relayed back via non-lemniscal thalamus as a second distal input to the apical dendrites of L2/3 and L5 PNs in L1, producing the MMN’s main negative deflection.

### Comparing animal and human neural dynamics with simulated LFP and CSD

Our model’s CSD pattern aligns somewhat with macaque data showing a superficial sink consistent with synaptic input to the distal dendrites of the L2/3 and L5 PNs. However, differences between species, recording methods, and anatomical targets must be considered. Laminar recordings in non-human primates typically sample A1, whereas MEG responses likely reflect activity across A1 and auditory association cortices. Supporting this, one macaque study found no differences in A1 between deviants and standards at human MMN latencies^40^, consistent with both our source localization and prior work showing that the human MMN arises in higher-order auditory regions on the anterior superior temporal plane. Thus, while SSA-related effects in L2/3 of A1 may contribute, projections to L5 in association cortex may be required for full MMN generation.

### Distinction from prior auditory modeling study of perceptual awareness

In previous work (Fernandez Pujol et al., 2023), we modeled detected target tones in noise, which elicit a P1-N1 complex and an auditory awareness negativity (AAN, ∼200 ms), with a similar proximal-distal-distal input sequence. Despite qualitative similarities, differences in timing, amplitude, and function distinguish the AAN from the MMN. The AAN reflects perceptual awareness, peaking later, while the MMN indexes deviance detection. Circuit mechanisms also differ: the AAN primarily involves distal input to L5 apical dendrites, whereas the MMN involves concurrent distal input to both L2/3 and L5. Although the origin of the AAN’s second distal input remains uncertain, evidence suggests the MMN’s second input arises from the non-lemniscal thalamus (Lakatos et al., 2020).

## Limitations

### True deviance detection versus SSA

Our stimulation paradigm lacked a “many-standards” control (Schröger and Wolff, 1996), preventing full dissociation of true deviance detection from release from SSA. Although we counterbalanced physical tone frequencies across trials, rarity effects may still have contributed to the deviant-standard difference (Schröger and Wolff, 1996; May and Tiitinen, 2010; Ross and Hamm, 2020). Thus, our findings may reflect a combination of true deviance detection and SSA-related mechanisms. Previous modeling work (Ross and Hamm, 2020) SSA increases input to SST interneurons (not explicitly modeled here), which inhibit PV cells, thereby disinhibiting L2/3 PNs. However, despite this disinhibition, reduced excitatory input due to SSA limits the generation of large, negative-going responses such as the MMN. Relatedly, we did not explicitly model potential synaptic depression effects in the first proximal input.

### Lack of specific inhibitory interneurons

HNN’s architecture includes only a single class of inhibitory neuron, functionally resembling PV cells, whereas real cortical circuits also include SST and vasoactive intestinal peptide (VIP) cells, with distinct laminar targets (Tremblay et al., 2016; Urban-Ciecko and Barth, 2016). The current model’s L2/3 inhibitory neurons generate both fast (GABA_A_) and slow (GABA_B_) inhibition onto L2/3 PN somata (and L5 PN apical dendrites). True L2/3 SST neurons predominantly target distal L2/3 PN dendrites, while L2/3 PVs target perisomatic compartments (Contreras et al., 2019). Thus, our inhibitory interneurons are effectively

PV-like. Although HNN does not explicitly model SST cells, we approximated their effects by modulating their synaptic targets. Future work incorporating multiple interneuron subtypes could refine our understanding of inhibitory contributions to MMN generation.

### Lack of neurons in L6

HNN’s architecture does not include L6 neurons, which may be important for both signal generation and modulation.

Though possessing smaller and less aligned dendritic trees than L5 PNs (Ledergerber and Larkum, 2010; Feldmeyer, 2023), L6 PNs nevertheless have relatively elongated apical dendrites (longer than L2/3 PNs, for example) and thus may still contribute to the MEG signal. Moreover, they regulate activity in superficial layers through intracolumnar and cortico-thalamo-cortical feedback loops (Sherman and Guillery, 1998; Dash et al., 2022). Two main L6 populations exist: (i) L6A CT neurons projecting to the lemniscal thalamus, and (ii) L6B neurons projecting to the non-specific matrix thalamus and within a column to apical dendrites of L5B PNs (Zolnik et al., 2020; Feldmeyer, 2023; Whyte et al., 2024).

L6A neurons may convey cortical predictions to the lemniscal thalamus, where they are compared with sensory input and relayed back to the cortex (Varela et al., 2024). This could account for frequency-dependent modulations in MMN amplitude (Näätänen et al., 1988; Tiitinen et al., 1994). However, because stronger L4 input would produce a positive-going dipole, additional mechanisms (see, e.g., Bortone et al., 2014; Voigts et al., 2020) may be required to explain the MMN’s polarity. L6B neurons, conversely, project via the non-lemniscal thalamus and intracolumnar connections to L5B PN apical dendrites, producing distal input consistent with the second distal input modeled here.

### Earlier deviance-related components

Deviance detection has been observed in earlier components like the Nb and P50 (Grimm and Escera, 2012; Dykstra et al., 2016), which were not analyzed here due to filter settings. Future work could model these earlier components and their potential relation to SSA. Several studies suggest that SSA may be independent of NMDA receptor function (Farley et al., 2010) and may instead depend on AMPA-mediated synaptic depression (Kudela et al., 2018). Notably, SSA appears to arise *de novo* in both lemniscal and non-lemniscal auditory pathways, rather than being inherited from subcortical structures (Ulanovsky et al., 2003; Khouri and Nelken, 2015). These observations suggest that SSA reflects a locally emergent cortical process that may share some mechanistic overlap with, but remain distinct from, deviance detection. Future in vivo studies are needed to clarify its mechanistic relationship to deviance detection.

## Supplementary Materials

### Alternative models

We tested two alternative models motivated by prior studies. (1) An alternative model that attempts to describe the difference between the response to the standards and deviants by changing only L2/3 parameters consisting only of L2/3 mechanisms and activity (Ross and Hamm, 2020) (Fig. S1 and Fig. S2). This model, while partially consistent with ours, was not sufficient to account for the MMN in our data. (2) An alternative model with increased exogenous excitatory input to the L5 cells. This model produced a better fit amplitude-wise but not latency-wise (Fig. S3 and S4). Both models use a proximal-distal input sequence to simulate the responses elicited by standard and deviant stimuli.

### Alternative Model 1

Starting with the model to standard tones as described above, we focused on adjusting parameters that reflect testing the three main aspects of prior models (Ross and Hamm, 2020). (i) weaker input from upstream synapses to the L2/3 PNs (a result of synaptic adaptation) (reflected by the distal input to the L2/3 PNs that has decreased NMDA conductance) and (ii) stronger excitatory input to the L2/3 SST interneurons (synaptic facilitation) / weaker excitatory input to the L2/3 PV interneurons (reflected by the absent distal input to the L2/3 basket cells which we assume to be PV interneurons). To model the response to the deviant tones we: (i) increased the excitatory input to the L2/3 PNs by increasing the conductance of the L2/3 NMDA receptors on these cells and (ii) increased the input to the L2/3 PV interneurons (Fig. S2). Because this model does not account for downstream effects, we did not introduce a second distal input to account for the large negativity at around 130 ms (the MMN) and kept the input to the L5 pyramidal and basket cells fixed.

### Alternative Model 2

We also attempted to model the response to the deviant tones by increasing the excitatory input to the L5 PNs, as compared to the standard tone simulation (Fig. S4). Although increasing said input did account for the amplitude of the MMN, the peak latency between the modeled and recorded responses to the deviants did not match. To account for this shift in peak latency we also had to increase the average start time of the distal input.

## Acknowledgments

This research was supported by the University of Miami (ARD, CFP, JB), the National Center for Advancing Translational Sciences of the National Institutes of Health under Award Numbers UL1TR002736 (ARD) and U24NS129945 (SRJ), a McKnight Doctoral Fellowship from the Florida Education Fund (CFP), and the University of Central Florida (ARD). We also thank Dr. Yunkai Zhu, Dr. James Bonaiuto, and Miguel De Jesus Silveira for their insightful comments and feedback.

## Availability of data and material

All data and code required to reproduce the findings in this paper are available at: https://osf.io/3zgvc/?view_only=dac53dbb39a9499ca85eb9a5f87b2f1a

## Authors’ contributions

CFP, JB, and ARD devised the project. ARD collected the data and supervised the modeling work. CFP and JB performed the simulations. CFP, JB, SRJ, RVT, and ARD contributed to the interpretation of the results. CFP, JB, SRJ, RVT and ARD wrote the manuscript, which all authors critically revised and approved.

## Supplementary Figures

**Figure S1.**
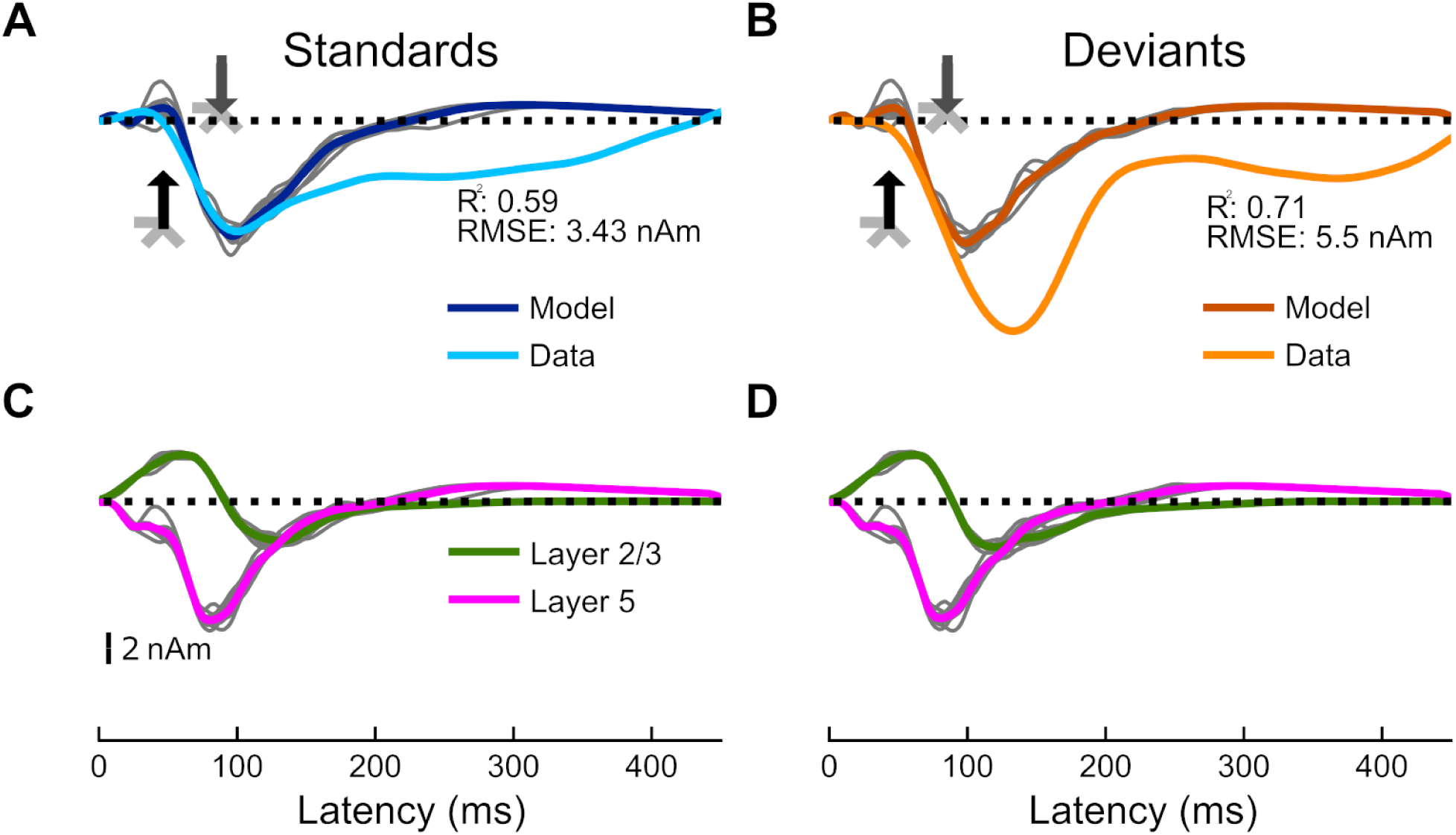
Current dipoles and laminar profiles for the first alternative model. (A, C) Current dipoles and laminar profiles for the main model presented here. (B, D) Current dipoles and laminar profiles for a 2-input (proximal and distal) model compatible with prior models (Ross and Hamm, 2020) for the response to the deviant tones. R^2^ between empirical (light orange trace) and simulated data (dark orange) is 0.71 and RMSE is 5.50 nAm. The corresponding input parameter values are displayed in Fig. S2.

**Figure S2.**
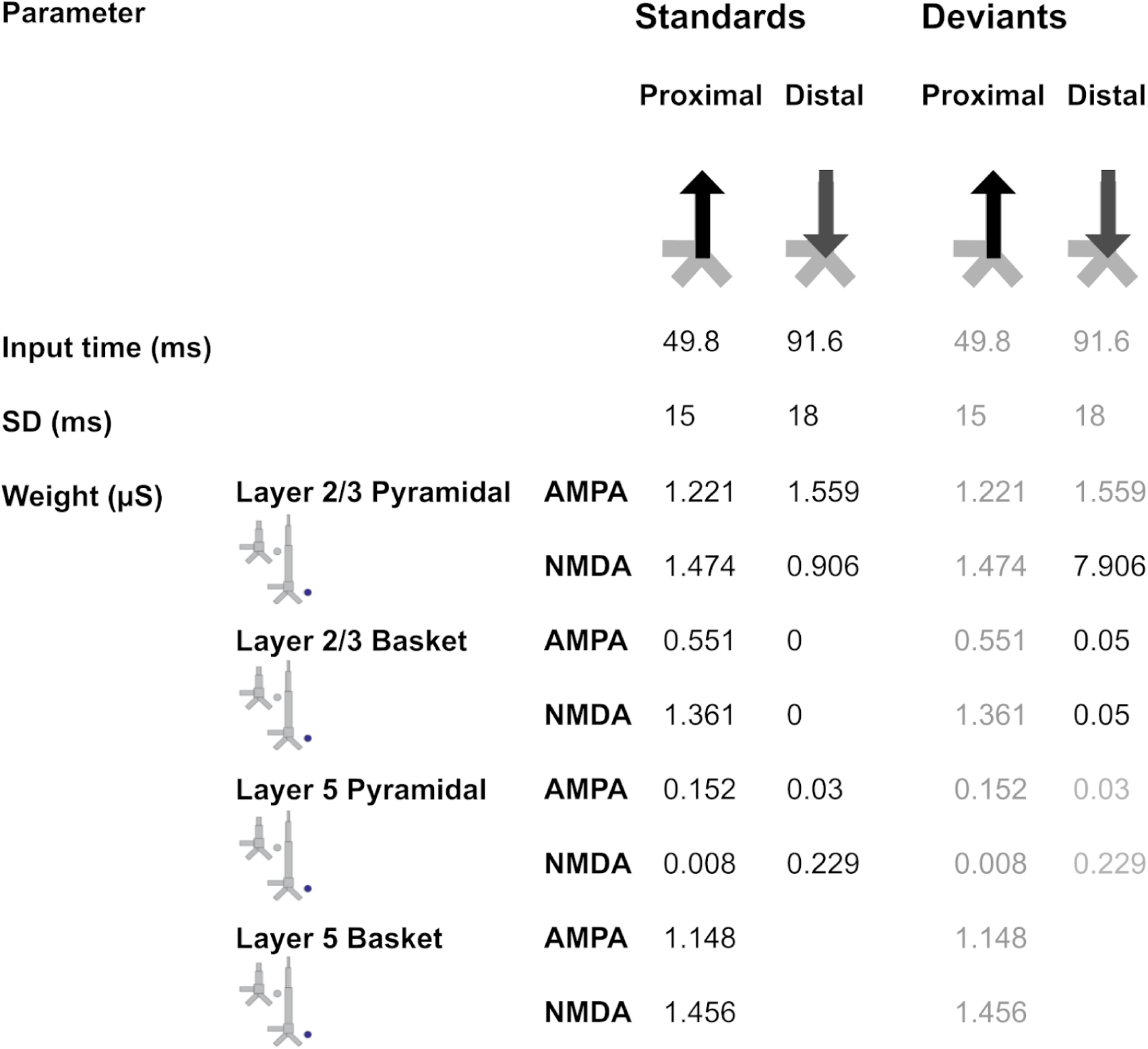
Driving inputs (and parameters) used to generate the first alternative model. The inputs and corresponding parameters used to model the standards are the same as those presented in the main model here (see Fig. 3). The parameters used to model the response to the deviant are consistent with models proposed previously (Ross and Hamm, 2020).

**Figure S3.**
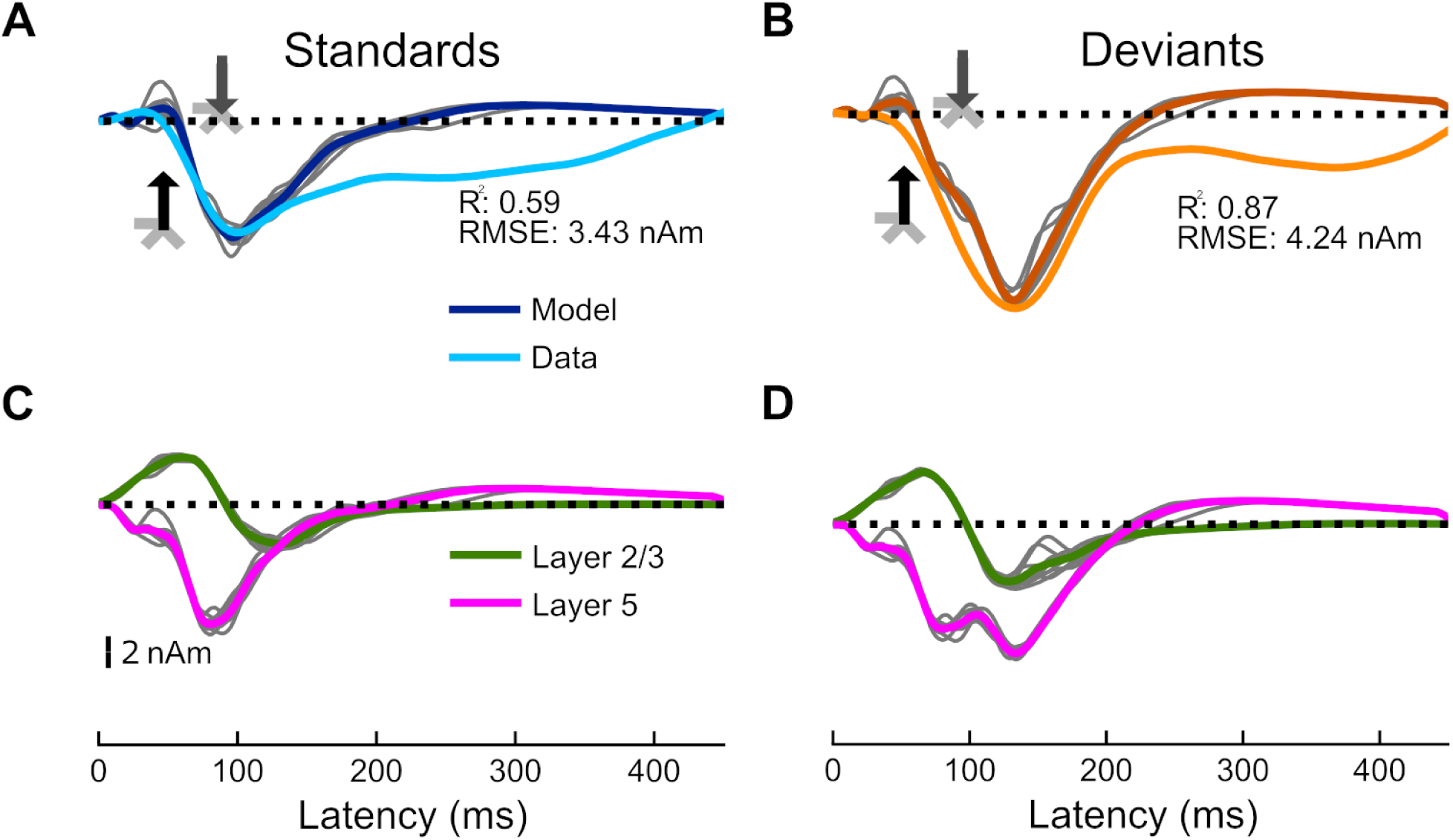
Current dipoles and laminar profiles for the second alternative model. (A, C) Current dipoles and laminar profiles for the main model presented here. (B, D) Current dipoles and laminar profiles for the same 2-input (proximal and distal) model as in Figs. S1 and S2, but with increased L5 PN conductances. R^2^ between empirical (light orange trace) and simulated data (dark orange) is 0.87 and RMSE is 4.24 nAm. The corresponding input parameter values are displayed in Fig. S4.

**Figure S4.**
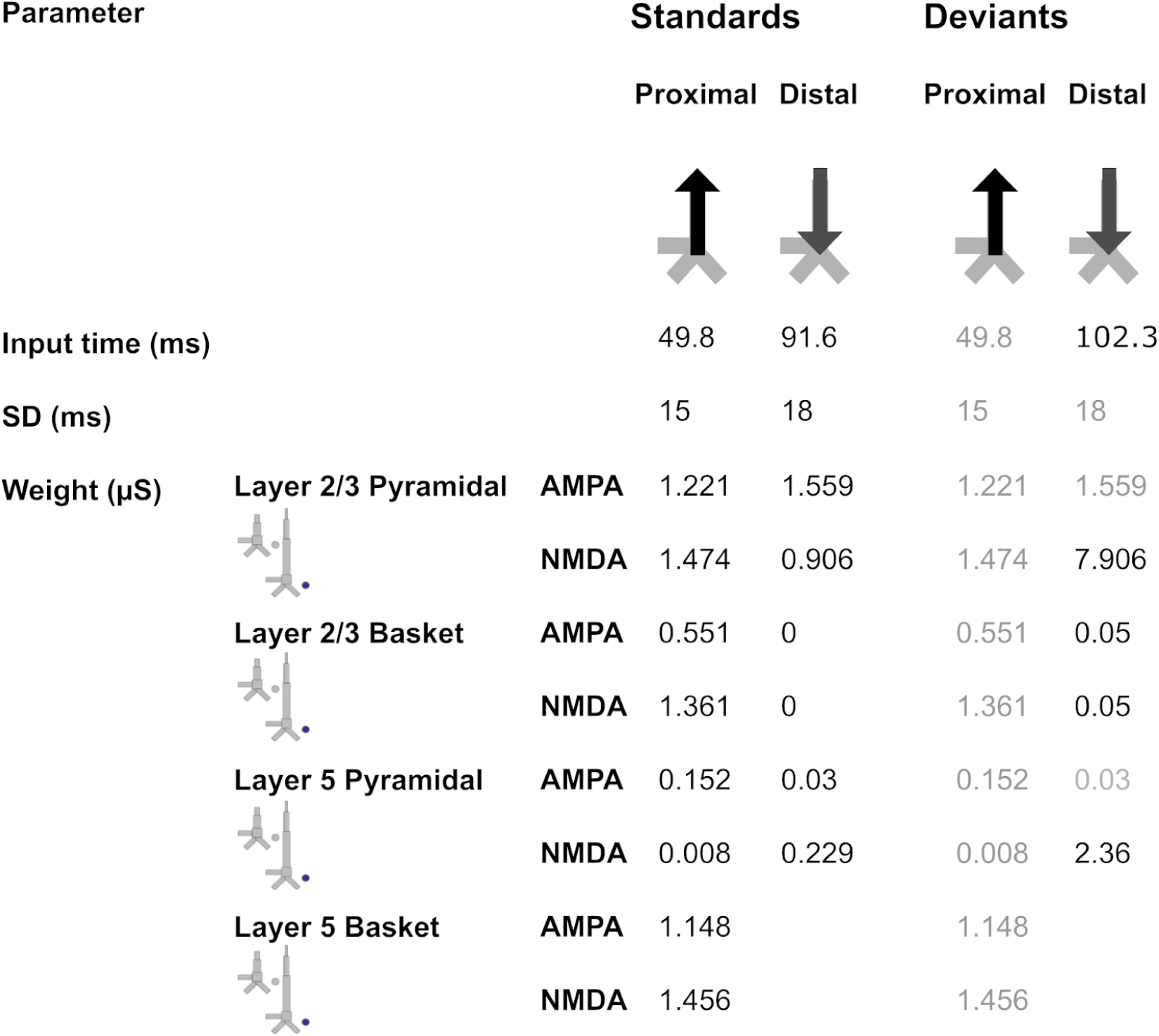
Driving inputs (and parameters) used to generate the second alternative models. The inputs and corresponding parameters used to model the standards are the same as those presented in the main model here (see Fig. 3). The parameters used to model the second deviant response correspond to the same parameters as in Fig. S2 (Ross and Hamm, 2020), but with increased input to layer 5.

